# Brood care in shell-dwelling cichlids is timed by independent maternal and larval clocks

**DOI:** 10.1101/2023.12.22.573114

**Authors:** Ash V Parker, Manuel Stemmer, Swantje Grätsch, Alessandro Dorigo, Oriolson Rodriguez Ramirez, Abdelrahman Adel, Alex Jordan, Herwig Baier

## Abstract

Successful brood care requires a finely tuned interplay between caregiving parent and offspring. Cichlids are a diverse group of teleost fish known for their extensive parental care of altricial young. Female *Lamprologus ocellatus* lay their eggs in the protective confines of an abandoned snail shell, where they raise them to free-swimming larvae. Here we investigated whether the transition to larval independence is driven by the behavior of the mother, the offspring, or an interaction between the two. Using 3D printed shells with a window, we were able to observe the behavior of both mother and offspring inside the nest using automated object detection and tracking. We found that, once hatched, the larvae remain in the deep chambers of the shell, and at 9 dpf, actively emerge from the shell during the day. An inversion of larval phototactic behavior from dark- to light-seeking leads up to this emergence. Removal of the mother causes premature emergence of the larvae by a few days. The natural schedule (emergence at 9 dpf) is restored if the larvae are supplied with fresh water in the mother’s absence. In cross-fostering experiments, we found that swapping out the mother’s own clutch for older larvae forces a delay in emergence, as the foster mother prevents the larvae from leaving at 9 dpf. We conclude that larval and maternal behavior are each controlled by independent internal clocks, which, although normally synchronized, can be experimentally manipulated and brought into conflict. This study has thus revealed an innate sequence of behavioral adaptations that orchestrate brood care in shell-dwelling cichlids.

## Introduction

Brood care requires synchronization of parental behavior with the developmental schedule of their offspring (Lock et al., 2004; Mousseau & Fox, 1998). Evolutionary forces have shaped distinct parental behaviors, including nest building, food provisioning, calling and guarding (Almada & Santos, 1995; Lock et al., 2004; Royle et al., 2012), while offspring have adapted through behaviors like begging, stress calls, and imprinting (Haack et al.,1983; Harper, 1986; Muller & Smith, 1978). For example in the burying beetle, *Nicrophorus vespilloides*, offspring beg for food, to which the parents respond by regurgitation (Lock et al., 2004). In songbirds, communication between parents and offspring is also required for “emergence”, i.e. the transition of altricial young from the nest to the outside world (Jones et al., 2020; Naef-Daenzer & Grüebler, 2016).

The timing of emergence is under strong selective pressure (Johnson et al., 2004; Jones et al., 2020; Naef-Daenzer & Grüebler, 2016; Nilsson & Svensson, 1993). Parents may benefit from their young leaving the nest early, reducing the time and expenditure needed until the arrival of the next brood (Johnson et al., 2017). However, extending the nest period can enhance offspring fitness. In songbirds, two hypotheses have been put forward: The ‘parent manipulation’ hypothesis, where parents and offspring are in conflict over the optimal fledging age (Jones et al., 2020; Martin et al., 2018), and the ‘nestling choice’ hypothesis, where fledging occurs when offspring reach a developmental milestone or when the costs of sibling competition outweighs the benefits of staying in the nest (Johnson et al., 2004, 2017; Nilsson & Svensson, 1993).

Extensive parental care is also found in some teleosts, particularly in the family *Cichlidae* (Balshine & Abate, 2021; Keenleyside, 1991; Sefc, 2011). Found mainly in East African Rift lakes, cichlids have evolved diverse brood care behaviors (Baerends & Baerends-Van Roon, 1950; Balshine & Abate, 2021; Sefc, 2011). Typically, embryos and larvae are endowed with yolk sacs, which provide nutrition for days to weeks (Kuwamura, 1997; Sefc, 2011). To safeguard their vulnerable offspring, cichlid parents either build nests or carry offspring in their mouth cavity (Sefc, 2011). As larvae reach a specific stage, they become free-swimming and venture out of these secure spaces.

Here we investigate the development and emergence of *Lamprologus ocellatus*, a shell-dwelling cichlid from Lake Tanganyika. These fish construct nests from abandoned gastropod shells (Koblmüller et al., 2007; Sato & Gashagaza, 1997, Balshine & Abate, 2021). *L. ocellatus* are obligate shell-dwellers, and reproduction starts with elaborate nest-building behavior (Baier, 2006; Lein & Jordan, 2021). Females lay the eggs inside the shell, where they are fertilized by the male, and raise the young fry inside the shell until they emerge. Even after emergence, *L. ocellatus* larvae return to the shell to sleep and when danger looms. After several weeks the now-juvenile fish move to the father’s shell for further protection (Bills, 1996; Haussknecht & Kuenzer, 1991). To disentangle the factors governing the emergence from the shell in *L. ocellatus*, we monitored the development of embryos and fry inside the shell. We discovered that emergence is timed by two intrinsic behavioral mechanisms, one of the mother, who prevents the larvae from leaving the nest until a prespecified stage, the other of the larvae whose phototactic behavior switches from dark to light preference in anticipation of emergence. Together, our study is the first to pinpoint innate mechanisms orchestrating maternal-offspring interactions in a teleost fish.

## Results

### Cichlid development and brood care can be monitored inside a 3D printed shell

*Lamprologus ocellatus* breed successfully in captivity, presenting behaviors similar to what has been reported in the lake, including egg laying and fry guarding (Fig. 1a,b,c,d) (Bills, 1996; Jordan et al., 2021; Sato & Gashagaza, 1997). We devised an artificial shell, modeled after an existing 3D model of the snail species *Neothauma tanganyicense* (Bose et al., 2020), and removed the back of the shell, opposite to the aperture, just behind the columella (inner spiral), and then extruded the cut walls outwards (Fig. 1e). This permitted the mother to navigate into the deeper chambers, replicating her natural behavior (Fig. 1f). We were also now able to virtually divide the shell into three compartments: entrance area, laying chamber, and deep chamber (Fig. 1g). The open-back shell was placed against the inside wall of the aquarium, with its open side facing the glass (Fig. 1h). Shells were provided for both the female and the male fish, and the inside of the mother’s shell was filmed for 11 days from the time of egg-laying, acquiring both timelapse snapshots and real-time video data in regular intervals (Fig. 1i). A near-infrared (NIR)-sensitive camera recorded the inside of the shell through an NIR-penetrable plastic (Fig. 1h). Embryos develop to larval fish inside the shell over this time period (Fig. 2a; Video 1). A machine learning object detector, YOLO v5 (Redmon et al., 2016), was trained to detect and locate the eggs and larvae across frames (Fig. 2b, 2c).

**Figure 1:**
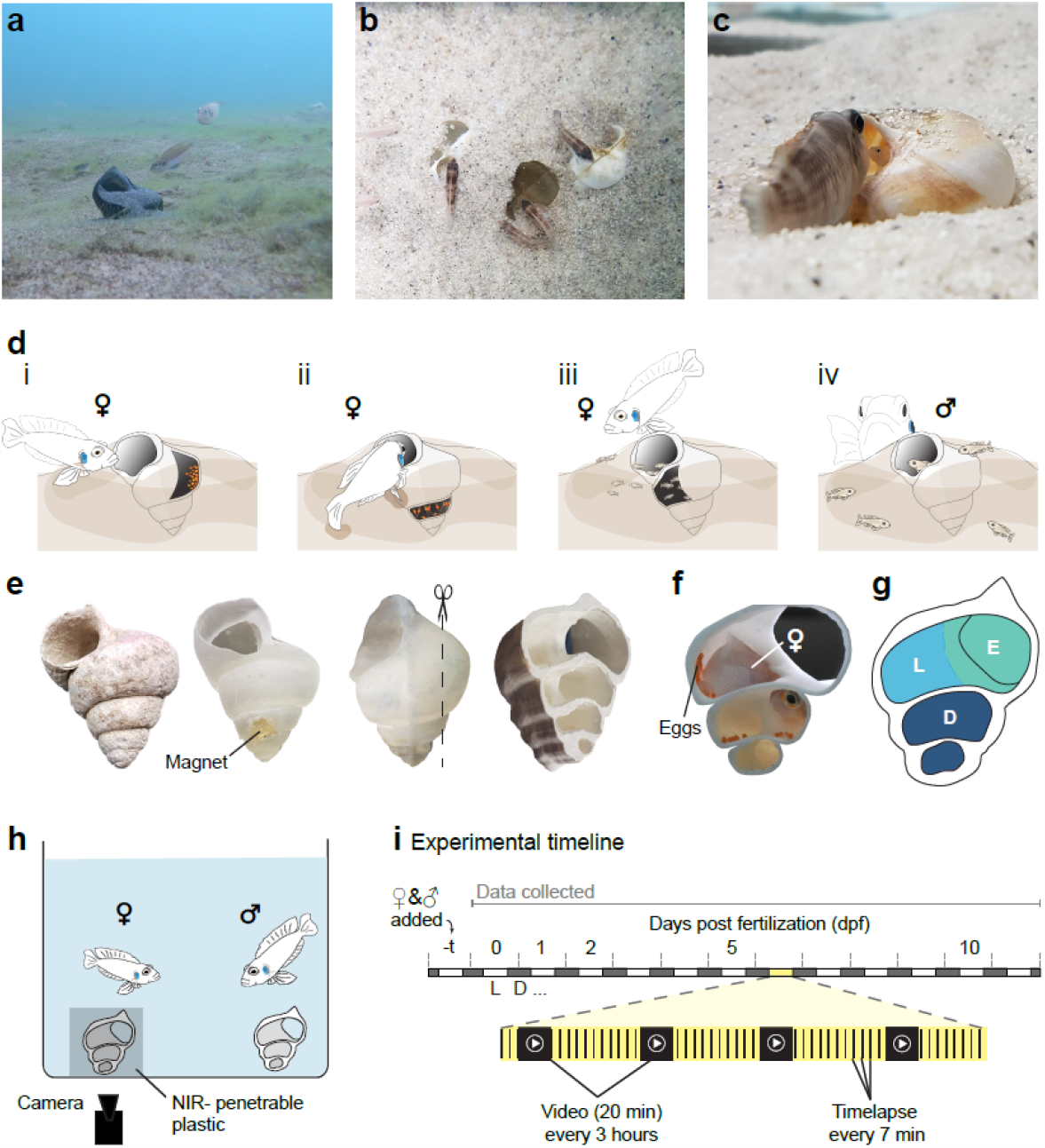
Observing *Lamprologus ocellatus* brood care inside an artificial snail shell. **a)** *L. ocellatus* male and female at the female’s shell buried in the sandy bottom of Lake Tanganyika (Photo credit Alex Jordan). **b)** *L. ocellatus* occupying and defending buried 3D printed shells in lab conditions. **c)** *L. ocellatus* father at his shell entrance with offspring inside the shell in our facility. **d)** Brood care stages of *L. ocellatus*: **di)** Mother laying eggs inside the first whorl of the shell nest, **dii)** hatched fry stay deep within the shell, **diii)** larvae are seen emerging from the shell, **div)** juveniles move across to the father’s shell. **e)** Construction of a 3D printable version of the Tanganyikan gastropod shell that allows camera footage access to the shell. **f)** Artificial extension of the dorso-ventral sides ensures access to deep chambers by adult *L. ocellatus* **g)** For analysis breakdown, the space inside the shell was divided into the entrance area (E; teal), laying chamber (L; blue) and deep chamber (D; navy). **h)** The tank set up includes a male and female breeding pair, each provided with a 3D printed shell attached to the glass and covered with a near infrared (NIR) -penetrable plastic to prevent visible light entering into the shell. **i)** The experimental timeline: Introduction of breeding pair occurs t days before fertilization. An NIR camera records images and videos from fertilization until the larvae are 11 days post-fertilisation (dpf) across daytime (L; white) and nighttime (D; gray).

**Figure 2:**
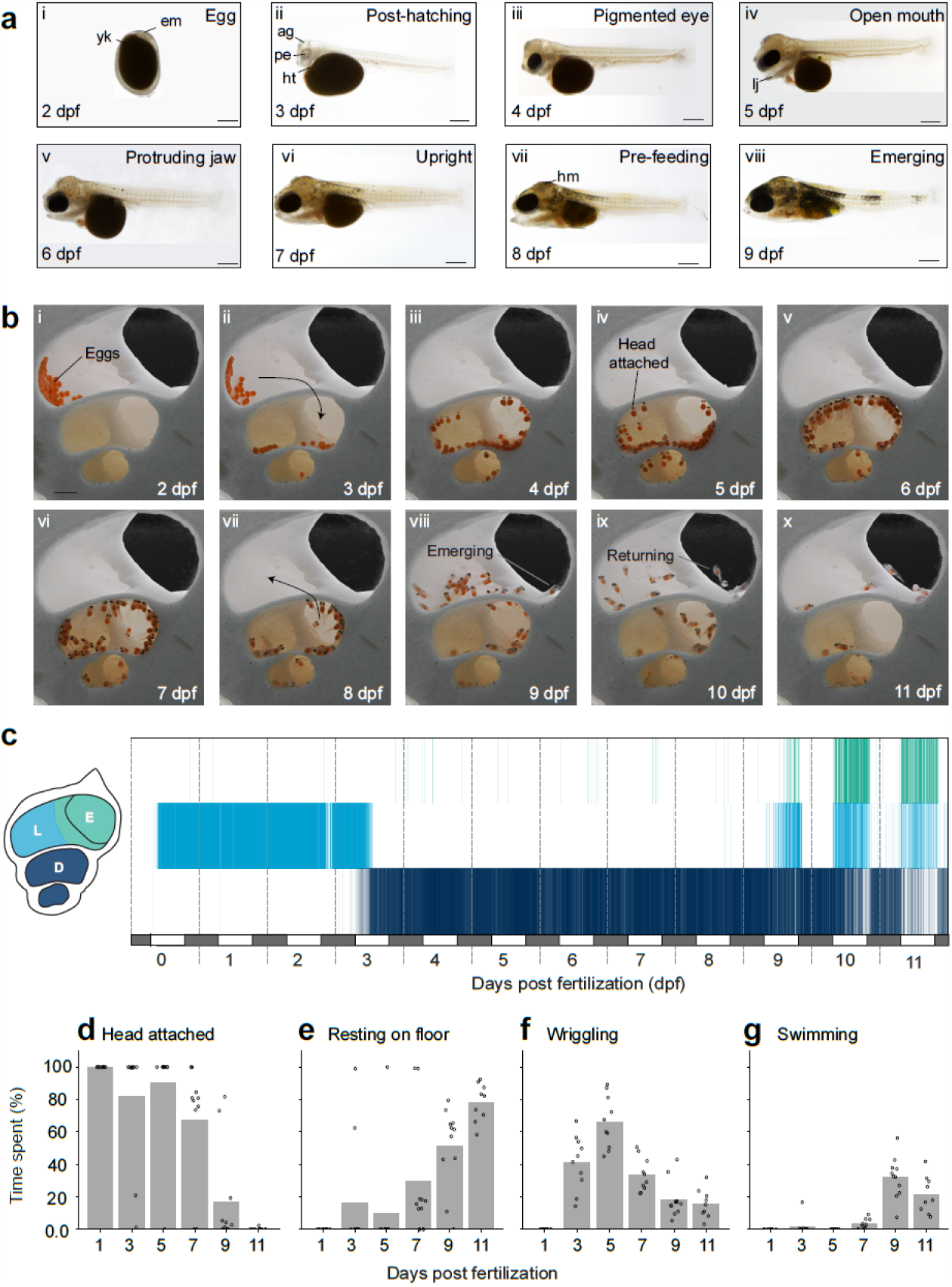
Larval morphological and behavioral development inside the shell. **a)** Morphological development: **ai)** Egg stage and **aii-viii)** Larval stages from 3-9 days post fertilization (dpf). Scale bar = 0.5mm. Abbreviations: yk, yolk; em, embryo; ag, adhesive gland; ht, heart; pe, pigmented eye; lj, lower jaw; hm, head melanophores. **b)** Shell space utilization: **bi**,**ii)** Egg stages, **bii-vii)** wriggler stages, and **bvii-x)** emerging larvae. Curved arrows represent larvae migration through the shell chambers. Scale bar = 5 mm **c)** Typical distribution of eggs and larvae in each chamber from 0-11 dpf from one clutch. The alpha value of each vertical coloured line in each chamber (E, entrance area; L, laying chamber; D, deep chamber) represents the normalized proportion of larvae at the time of the snapshot. White and gray alternating bars under the plot indicate day and night, while vertical dotted gray lines separate each dpf. **d-g)** Larvae (n=10) behaviors inside the shell: **d)** head attached; **e)** resting on the floor; **f)** wriggling; **g)** swimming. Larvae were observed for 5 minutes in video data and scored using BORIS (see Ext data table 1 for behavioral descriptions).

### Embryonic and larval adaptations help survival inside the shell

Females lay their eggs inside the shell, typically on the side walls of the first whorl (Fig. 2bi,c). Almost all egg-laying activity was observed in the morning, and took 1.99 ± 1.6 hours (mean ± standard deviation (sd), n=12). Clutch size ranged from 10 to 50 eggs with an average of 28±10 (± sd, n=20) and egg size averaging 1.42 ± 0.08 mm (± sd; n=114). The large orange yolk of cichlid eggs leaves minimal space for the embryo to develop within the sticky chorion (Fig. 2ai) (Kratochwil et al., 2015). After hatching, at 2 or 3 days post-fertilization (dpf), the mother transports the larvae into the deep chamber of the shell in her mouth, where they reside for the next 6 days (Fig. 2bii,c). Soon after hatching, an adhesive gland forms on the top of the larvae’s heads, a feature present in most substrate-brooding cichlid larvae (Balshine & Abate, 2021; Kratochwil et al., 2015; Peters & Berns, 1983). This gland serves to anchor the larvae to the shell wall before they become free swimming (Fig. 2aii-vi,biii-vii). At 3 dpf, the pigments of the eye are starting to appear, and the first red blood cells begin to circulate from the heart (Fig. 2aii). In the shell, the larvae can be found either attached to the shell or resting on the shell floor, as the density of their yolk weighs them down (Fig. 2bii-vii,d,e; Video 2). By 4 dpf, the eyes are fully pigmented and the head and body have grown. The lower jaw is also now visible (Fig. 2aiii), becoming prominent at 5 dpf (Fig. 2aiv). At 6 dpf, the lower jaw becomes free from the yolk, extending the head in an anterior direction (Fig. 2av). From this stage, the larvae are constantly opening and closing their mouths. Melanophores start appearing on the head and dorsal side of the larvae. At 8 dpf, the melanophores increase in density on the head and anterior spine (Fig. 2avii). By 9 dpf, an additional two patches of melanophores appear along the spine, alternating with areas lacking pigment, giving the larvae a striped appearance (Fig. 2aviii).

Once hatched, larvae perform intervals of high-frequency tail beating, termed “wriggling” (Video 2; Courtenay & Keenleyside, 1983). Wriggling bouts may last from 0.03 to 50 seconds. The amount of time spent wriggling varies as the larvae age (Fig. 2f). At 3 dpf, larvae are head-attached 82% of the time and spend an average of 41% of the time wriggling. At 5 dpf, larvae are still attached to the walls of the shell by their head glands, but their time spent wriggling reaches its peak at an average of 66% (Fig. 2d,f). At 7 dpf, the time spent wriggling decreases back to an average of 34% and while larvae are still head-attached 68% of the time, 29% of the time they are lying on the shell floor. Some larvae were recorded detaching (‘swimming’) for a maximum of 6.7 seconds before again attaching to the shell, or resting on the floor (Fig. 2 b,d-g; Video 2). Notably, the larvae resting on the floor undergo a shift in posture between 7 and 8 dpf, from lying on their sides to ‘upright’ (ventral-side down; Fig. 2avi,bvii), possibly as a result of inflation of the swim bladder (Kratochwil et al., 2015). At this stage, larvae were exclusively observed to reside in the deep chambers (Fig. 2c).

### Emergence from the shell follows active exploratory movements of larvae at 9 dpf

At 9 dpf, the larvae undergo an important transition, reproduced in all of our control observations: They actively move up to the laying chamber and into the entrance area (Fig. 2bix,c). There is a further decrease in wriggling duration to 20% of the time, and this coincides with an increase in swimming periods (Fig. 2f,g). The time spent head-attached decreases to 18%, and the time spent resting on the shell floor increases to 51% (Fig. 2d,e). We tested their ability to catch prey, by transferring larvae as young as 7 dpf to petri dishes and adding live brine shrimp (*Artemia salina*). Larvae only begin hunting this prey from 9 dpf onward, at which point we deem them independently feeding (see Mearns, Hunt et al., 2023). The last remnants of yolk remain noticeable until 11 dpf (Fig. 2bx).

From 9 dpf onward, behavior during the night becomes distinct from that of the day. During the day, the larvae enter and exit the shell, utilizing all chambers, whereas, at night, larvae are residing in the deep chamber (Fig. 2c). Over the ensuing days, larvae spend more time out of the shell during the day and use the deep chamber of the shell only at night for refuge (Fig. 2c).

### Before emergence from the shell, larval behavior switches from dark to light preference

We asked what larval-intrinsic factors drive the emergence from the shell. To determine if emergence is caused by hunger, we fed the larvae inside the shell with a thin tube going directly into the deep chamber. Small quantities of live brine shrimp were syringed in the shell from early 9 dpf throughout the day. Video data showed that the larvae did perform prey capture within the shell and fed on brine shrimp, but this did not delay the emergence from the shell (Ext. data Fig. 1, n=1). This observation suggests that motivation to search for food contributes little, if any, to leaving the nest.

We next tested if pre-emergent and emergent larvae were attracted to light and therefore migrated out of the shell during the day. Larvae were collected from shells at 6 dpf and placed in a phototaxis testing arena in small groups of 4 to 5 animals (Fig. 3a). One half of the arena was dark and the other half exposed to the same ambient light cycle as in our previously described experiments (see Methods for details). Sides were switched daily. Images of the experiment were collected multiple times a minute over four consecutive days (7-10 dpf), and YOLO v5 was trained to detect larval positions inside the arena. The experiment shows that, at 7 dpf, almost all larvae were in the dark both day and night (Fig. 3b, n=28). At 8 dpf, on average 72% of the larvae still prefer to stay in the dark chamber, but this decreases as the day proceeds to an average of 65% in the dark. At 9 dpf, the preference tilts to migrating to the light: Only 40% of the larvae remain in the dark during the day. Similar averages are seen at 10 dpf. At night, when ambient light drastically decreases, there is a persistent preference for the dark across all ages observed, with averages between 56-100% of larvae on the dark side of the arena. Overall there is a significant decrease (p=0.03, two sided Wilcoxon signed-rank test) in daytime dark preference of pre-emergent stages (7 and 8 dpf), with an average of 76% in the dark compared to post-emergence (9 and 10 dpf) with an average of 38% in the dark (Fig. 3c).

**Figure 3:**
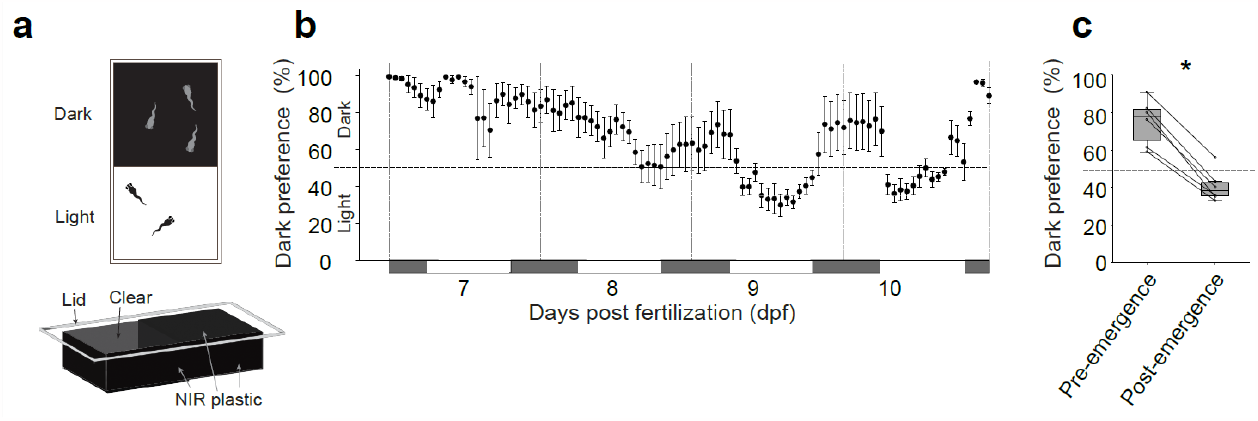
Switch in larval light preference corresponds to emergence time. **a)** The experimental box used to test light preference with one side of the lid blocking visible light (black rectangle, NIR plastic) and the other allowing visible light through (white rectangle, clear). **b)** The mean percentage and standard error of larvae (n=28) that preferred the dark side of the box in each hour across 7-10 dpf. White and gray alternating bars under the plot indicate day and night, while vertical dotted gray lines separate each dpf **c)** A box plot of the percentage of larvae in the dark during the daytime before and after emergence. Each circle indicates the mean preference in each experiment before and after emergence. Overall difference is significant (asterisk) (p=0.03, Wilcoxon signed-rank test).

### Maternal behaviors serve to maintain water quality and hygiene of larvae inside the shell

The mother performs active brood care behavior during the observation period. We used the YOLO object detection tool, trained with a model specifically for the mother’s head, and recorded her location within the shell (Fig. 4a). From this data we could measure visitation frequency over the 11 days (Fig. 4b, n=4-12) and also compare daytime and nighttime presence in the shell (Fig. 4b,c). The mother mostly visits the entrance of the shell (Fig. 4a), although she can reach the deep chamber with apparently little effort. At night the mother is almost constantly present in the entrance (Fig. 4a,c). During the day, the mother visits all three chambers (Fig. 4a). At 2 and 3 dpf (Fig. 4a,b), the mother’s visitation peaks, compared with other daytime periods, which coincides with larval hatching (Fig. 2b).

**Figure 4:**
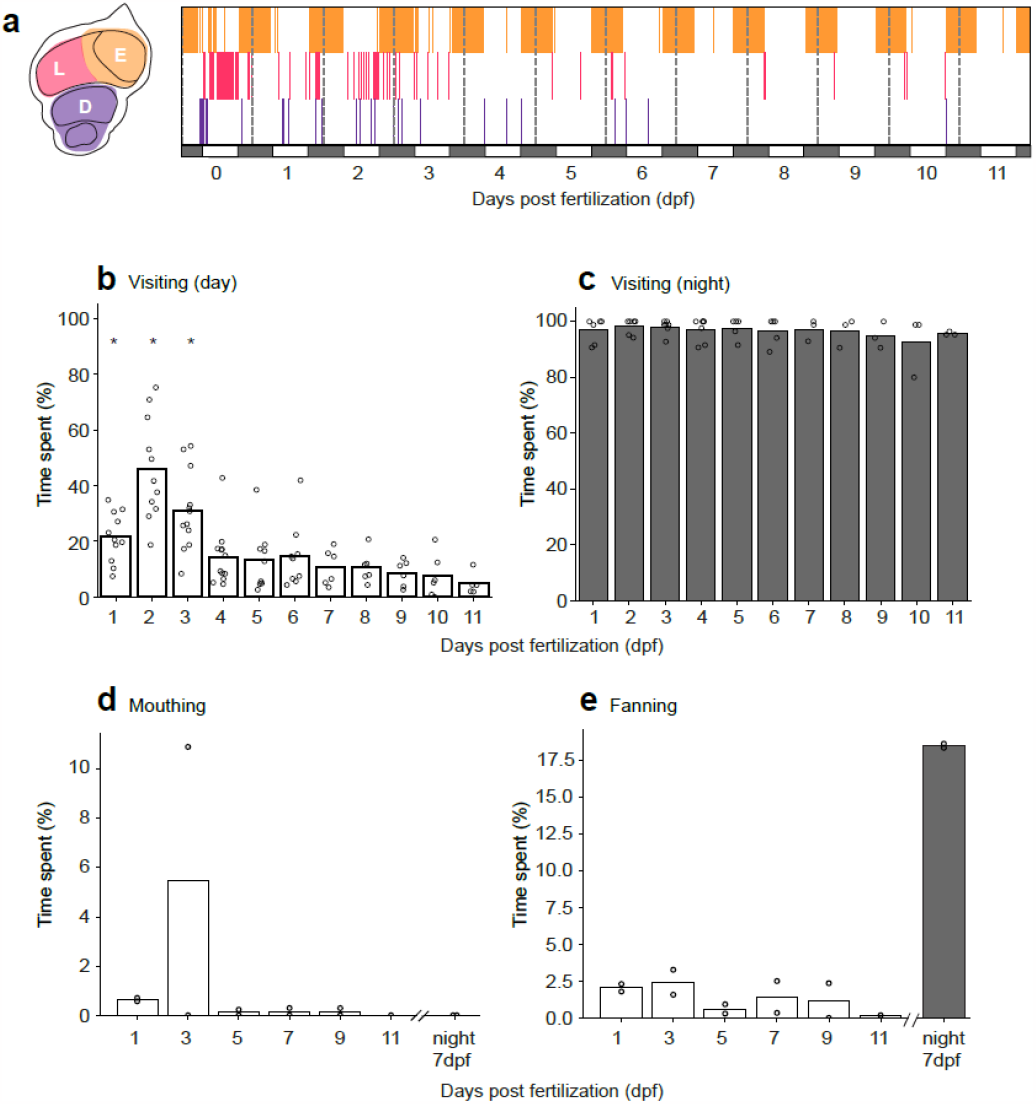
Mother’s behavioral repertoire inside the shell during brood care. **a)** The mother’s location inside the shell during the first 11 days of brood care. Automated mother detection (YOLO) and positional classification were used. Each vertical coloured line indicates the presence of the mother’s head in one of the chambers (E, orange, entrance; L, pink, laying; D, purple, deep) for each snapshot taken. White and black alternating bars under the plot indicate day and night, while vertical dotted gray lines separate each day post fertilization (dpf). **b&c)** The percentage time mothers spend in the shell during the **b)** daytime (n=4-12) and **c)** nighttime (n=3-5) from fertilization until 11 dpf. Bars graph represents the mean percentage across mothers recorded for that day/ night and circles are the values for the individual mothers. **d&e)** The percentage time mothers spend mouthing (n=2) **(d)** or fanning (n=2**) (e)** the eggs or larvae during 20 min video recordings at noon, daily or 3 am at 7 dpf. Bars graph represents the mean percentage across mothers recorded at each dpf and circles are the values for the individual mothers.

With assistance from the software BORIS (Friard & Gamba, 2016), the mother’s behaviors inside the shell were scored during 20 minute videos, taken around noon at 1, 3, 5, 7, 9 and 11 dpf and at 3 am at 7 dpf for nighttime behavior (Fig. 4e,f, n=2). Two key behaviors were noted in the shell (Video 3). The first was mouthing: Here the mother is seen either nipping at, or placing her lips on, the eggs to clean them and remove dead eggs. During hatching, mouthing becomes more frequent (Fig. 4d). Mothers pick up the hatchlings in their mouth and perform buccal cleaning with the offspring before depositing them down into the deep chamber of the shell (Fig. 2bii). The second behavior is fanning, where the mother rapidly moves her anal and caudal fin to facilitate water exchange inside the shell. Fanning behavior during the day is seen fairly consistently over the time the larvae are confined to the shell (0-9 dpf), with a slight decrease at 5 dpf (Fig. 4e). Very little fanning occurs at 11 dpf. At night (3 am at 7 dpf), fanning occurs approximately 20 times more than during the day (Fig. 4e).

### Emergence of orphaned larvae depends on provision of fresh, oxygenated water

In order to determine the maternal contribution to the emergence time of the larvae, we removed the parents. When the mother was removed pre-hatching, none of the eggs survived. Removal at either 5 or 7 dpf led to the larvae immediately moving up into the laying chamber (Fig 5b,f; n=5). The larvae then began to use the entrance chamber and emerged within 24 hours of the mother’s removal, premature in comparison to the control conditions (Fig. 5a, f; n=4). After emergence, day-night behavioral differences were less pronounced compared to the control where larvae reside exclusively in the deep chamber at night, but explore all three chambers during the day. In all these experiments, by 11 dpf, none of the orphaned larvae returned to the shell anymore.

**Figure 5:**
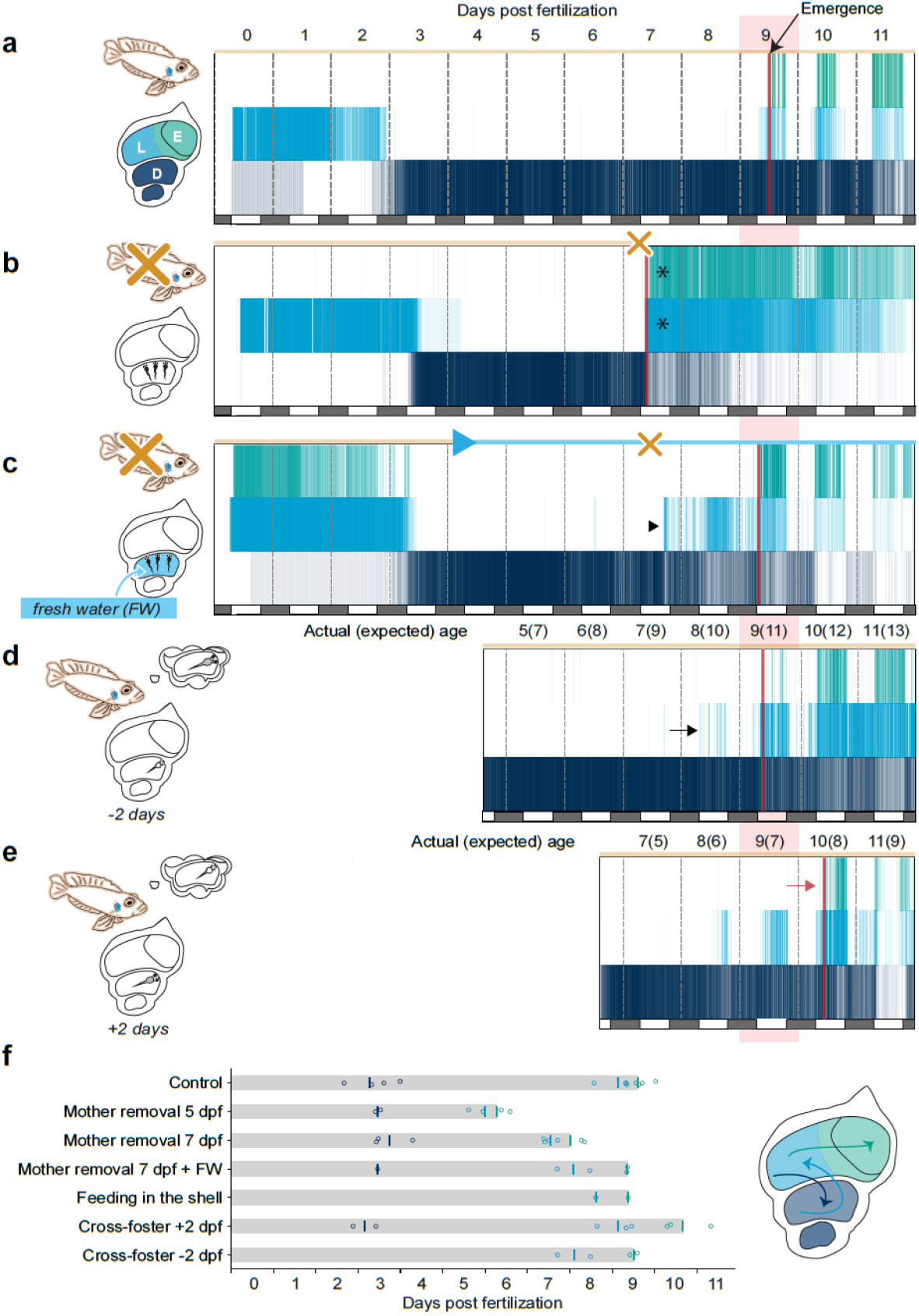
Larval distribution and emergence adjustments during brood care manipulations. Larvae distribution across three chambers (E, teal, entrance; L, blue, laying; D, navy, deep) over 11 days post fertilization (dpf) with presence of the mother (orange bar) and emergence time indicated (red line). Each figure is an example of the **a)** control condition, **b)** removal of the mother at 7 dpf (orange X), **c)** removal of the mother at 7 dpf while supplying larvae with continuous fresh water from 4 dpf (blue triangle and blue bar), and two cross-foster experiments of larvae **d)** 2 days younger than mother’s original clutch and **e)** 2 days older than mother’s original clutch. White and gray alternating bars under the plots indicate day and night, while vertical dotted gray lines separate each day. Expected emergence on day 9 is indicated by a transparent red bar behind the plots. **f)** Average transition time to the deep chamber (navy), then to the laying chamber (blue) and average emergence time (teal) of larvae across different manipulations, individual points correspond to each experiment.

In the absence of the mother, emergence happened more quickly under crowded conditions. In two experiments, in which clutches were 40 or 49 larvae, respectively, larvae were observed outside the shell within an hour after the parents’ removal at 5 or 7 dpf. Larvae from clutches of 24-34 larvae emerged substantially later. We hypothesized that deteriorating water quality inside the shell drives out orphaned larvae prematurely. To test this, we removed the parents again at 7 dpf, but supplied the larvae with a source of fresh water continuously from 4 dpf directly into the deep chamber of the shell. Again, the larvae moved into the laying chamber earlier than seen in the control condition (Fig 5c,f; n=2). However, larvae were not seen emerging before the expected 9 dpf. Additionally, the emergent larvae initially maintained a day-night behavioral rhythm similar to the controls by returning to the deep chamber at night, suggesting that oxygenation and/or waste removal by the mother are key factors in keeping emergence on schedule.

### Cross-fostering experiments show that the mother has an independent clock

The mother’s brood care behavior might follow an independent timer, e. g., started by the hormonally salient act of egg laying. Alternatively, she might update her behavioral responses on an ongoing basis by assessing the signals and status of her offspring. To decide between these possibilities, we switched the clutches of mothers for offspring 2 days younger or 2 days older than her original clutch and observed both maternal and offspring behaviors.

In all five of six experiments, the foster mothers accepted the foreign offspring. In experiments where mothers adopted offspring 2 days younger than expected by her, the larvae moved up to the laying chamber earlier than sibling controls that had stayed with the original mother and emerged according to their biological age at 9 dpf (Fig. 5d,f; n=2). In contrast, in the experiments where mothers fostered offspring 2 days older than expected by her, emergence was delayed by at least one day (at 10 or 11 dpf; Fig. 5e,f; n=3). Data on the mother show that she performs more mouthing and fanning than typically seen on this day in comparison to controls (Ext. data Fig. 2). Video data show the mother vigorously fanning larvae, creating a strong inward current, pushing the clutch deeper into the shell, and even picking up stray larvae from the laying chamber (Video 4). Together, these experiments indicate that the mother provides brood care on an intrinsically timed schedule.

## Discussion

In this study, we filmed, for the first time, embryonic and larval development of the shell-dwelling cichlid species *L. ocellatus* inside the protective confines of their nest. By using artificial neural networks, we tracked the distribution of fry inside the shell and recorded the parent-offspring interactions over an 11-day period from the time of fertilization (day 0) to a late-larval stage. Larvae emerge from the shell at 9 dpf and afterwards return to it only to hide or to sleep. This novel experimental paradigm has allowed us to discover key factors contributing to the larva’s decision to leave its nest, including internal timing mechanisms (clocks), which coordinate the mother’s and the offspring’s behavior in and around the shell.

Several notable behavioral and morphological adaptations of brood care in *L. ocellatus* correspond to the specific selective pressures of the shell environment: maintaining quality and oxygen content of the water inside the shell, providing hygiene, and avoiding predation. We observed that the mother regularly cleans the eggs and larvae likely from mold and bacteria by taking them up in her mouth, swishing them through her buccal cavity, and spitting them out. After hatching, at 2-3 dpf, she moves them to the deepest chamber and actively prevents them from leaving the shell prematurely. She may even pick up a stray hatchling, which has managed to slip out, and return it to the shell. She also vigorously fans the brood with her caudal fin pairs. Finally, after the larvae have emerged from the nest and swim freely nearby in search of food, she still signals potential threats and keeps guarding the shell entrance when the larvae are inside (e. g., at night and when danger looms). We found that key trends in maternal behavior follow a rigid schedule, which appears to be controlled by an intrinsic, perhaps neuroendocrine, clock (see, e. g., Bender et al., 2008). Heterochronic cross-fostering experiments showed that this maternal clock cannot be reset by exposure to either younger or older fry. Although she eagerly adopts a stranger’s offspring, an *L. ocellatus* foster mother will treat the fry as if they were her original clutch’s age and tries to ‘override’ the larvae’s schedule by blocking their emergence. Other cichlids, notably *Cichlasoma citrinellum* and *Hemichromis bimaculatus*, have been observed to extend their parental care when their brood was repeatedly replaced by younger fry (Noakes & Barlowy, 1973; Noble & Curtis, 1939). This earlier work suggests that the maternal clock may be more or less adaptable across different cichlid species.

*L. ocellatus* fry possess their own set of adaptations to growing up inside a shell. As embryos, they are small and loaded with nutritious yolk, which lasts well into the late-larval stage (11 dpf). An adhesive gland on their head and the delayed formation of a swim bladder constrain their motility until shortly before emergence (Balshine & Abate, 2021). After hatching, they show a whole-body shaking behavior (‘wriggling’), which is predicted to aid in water exchange for physical self-cleaning and oxygen enrichment (Baerends & Baerends-Van Roon, 1950; Yoshida et al., 1996). When deprived of fresh water (e. g., following removal of the mother), larvae leave the nest on an accelerated schedule, suggesting that (i) they sense oxygen (or another water quality parameter) and that (ii) low water quality can override their intrinsic schedule.

Strikingly, the phototactic behavior of *L. ocellatus* undergoes a precisely timed sign inversion when the larvae are ready to emerge: While young larvae in the first week of development swim away from the light (which under normal conditions would drive them deep into shell), their behavior switches from negative to positive phototaxis shortly before they leave the nest. A similar dark-to-light inversion of phototaxis has been observed in starling nestlings (Minot, 1988). In zebrafish, *Danio rerio*, such a developmental switch also occurs, but is reversed; i. e., from light-seeking in larvae to dark-seeking in juveniles (Lau et al., 2011). Interestingly, older *L. ocellatus* larvae still preferred darker places at night, i. e., migrate into the deep parts of the shell, a behavior that could act to protect them against nocturnal predators. A similar dark-seeking behavior was noted in jewel fish larvae when scared (Noble & Curtis, 1939). Feeding larvae with brine shrimp inside the shell before or around the time of emergence did not delay, or reduce, outside excursions. This suggests that hunger has little, if any, effect on the timing of emergence. It would be worthwhile to explore the effects of oxygen deprivation on phototaxis of *L. ocellatus* larvae, as a decrease in water quality at 7 dpf can apparently override the preference for darkness.

The two clocks, the maternal one and the larval one, are normally synchronized, resulting in a reproducible emergence time at 9 days under our laboratory conditions. At 2 weeks, when the fry are more independent and their mother is ready to raise the next clutch, she evicts them from her shell, whereupon they move in with their father into his shell (Haussknecht & Kuenzer, 1991). This complex series of interactions may be the echo of an ancient parental-filial conflict about the proper emergence time. Evolutionary theory predicts that the larvae should seek to extend their stay in the nest to maximize resource allocation afforded by the mother’s care and protection, while the parents are interested in keeping brood care obligations to the necessary and sufficient minimum to maximize the number of reproductive cycles (Jones et al., 2020; Mainwaring, 2016). Interestingly, the mother could, but does not, cut the residence time short by stopping fanning, suggesting that, in the case of *L. ocellatus*, ‘maternal choice’ and ‘nestling choice’ have found a stable equilibrium by genetically setting independent internal clocks in both parents and offspring. In this co-adaptation scenario, the timing of the phototactic switch in larvae is synchronized to the gradual, likely hormonally driven, cessation of maternal care.

## Methods

### Animal experiments

*Lamprologus ocellatus* were bred in 51 liter tanks (600 × 270 mm, 320 mm deep) with one male and two females at the Max Planck for Biological Intelligence, Martinsried, Germany, following institutional guidelines set by the Max Planck Society and regulations by the regional government (Regierung von Oberbayern). The colonies originated from wild-caught populations collected at Isanga Bay, Zambia. Tanks were filled with 4-5 cm of beach sand (0.4-1.4 mm grain-size) and for each individual one of two types of 3D printed Tanganyika shell was provided: the males typically claim the larger complete shell and the females took up one of two open-back shells, which were pressed up against the glass and protected from stray light by opaque tapes, allowing daily checks for the presence of new egg clutches (models provided and adapted from (Bose et al., 2020). Breeding fish, as well as fish during all experimentation, were fed live artemia daily and kept under constant conditions (13-11h light-dark cycle, 27°C water, pH 8.2, conductivity ∼550 μS). Between 7 am and 8 pm the room lights were on while at night these all went off, leaving only a faint light from aquarium computer monitors.

### Staging embryonic and larval development

Freshly fertilized eggs were collected in a 100 mm petri dish from the shell of a mother in the breeding tanks and the larvae were raised at 20 fish/ dish in a 27°C incubator with a 14 h - 10 h light-dark cycle, with fresh facility water changed daily. Eggs and larvae were visualized under a stereo microscope (Nikon SMZ25) with a 1x SHR Plan Apo objective. During brightfield image acquisition the exposure time and illumination intensity were adjusted using the NIS-Element imaging software.

### Behavioral observations inside the shell

An established breeding couple was introduced into a custom-built 100 liter experimental tank (420 × 660 mm, 360 mm deep) (Ext. data Fig. 3), with light-absorbing white acrylic bottom and sides and into which two 3D printed shells (stl file provided), 430 mm apart, were inserted to mimic a typical shell successfully buried into the sand, as no sand was provided. On the outside of the tank, the open shell-backs were protected from visible light with NIR-penetrable plastic covers (LUXACRYL-IR,1698, 0.8 mm thick, Germany). The tank was illuminated constantly with six 850 nm near-infrared light strips (NIR; 12 Osram-Olson LEDs, 700ma) from above and two 850 nm NIR light strips directed from the side into the filmed shell. This tank was part of the housing facility, with constant water exchange and visible lighting, water conditions and feeding were as described above. A NIR camera (Ximea, MX042RG-CM-X2G2-FF, Germany) with a 50 mm lens (Edmund Optics, Germany) was directed into the shell back, with the shell-back taking up the whole field of view and the aperture and focus adjusted to ensure the greatest focusing depth within the shell. Media acquisition and processing was programmed with custom-written python scripts and Ximea software to acquire images (2048×2048 pixels; JPEG) every 7 minutes and take 20 minute videos (30 frames per second, avi) at specific times of the day from fertilization of the eggs until and including 11 days post fertilization (dpf).

To assess the behavioral repertoire of the larvae and mother inside the shell, we annotated behaviors with the assistance of BORIS software (Friard & Gamba, 2016)). For the larvae, 10 individuals were observed and scored for 5 minutes at noon on 1, 3, 5, 7, 9 and 11 dpf across two independent control experiments. Behaviors annotated included visible, head attached, resting on floor, wriggling and swimming (see Ext. data table 1 for behavior descriptions). For the mothers, individuals were scored over 20 minute videos at noon on 1,3,5,7,9 and 11 dpf as well as 3 am on 7 dpf. Behaviors scored included visible, fanning and mouthing (see Ext. data table 2 for behavior descriptions). Mothers across all experiments and dpf were scored when present, however certain dpf were excluded if there was any disruption to the natural behaviors such as introducing water flow or food to the shell.

Using the BORIS software we were able to export a csv file of the aggregated events of the mother and the larvae separately for each dpf and experimental run. We then used custom-written python scripts to process the data and produce figure 2d-g and figure 4d-e. Data for the mother and larvae were processed separately but for both we first concatenated all the results across experiments for each dpf, ensuring each individual was named uniquely. We then calculated the total duration each individual engaged in each behavior and normalized this by dividing by the total duration the individual could be seen (visible) in the time observed. The mean and standard deviation was calculated across all individuals for each dpf and behavior.

### Localization of mother and larvae inside the shell

To detect the mother and egg or larvae inside the shell, we used YOLO v5 and trained using a custom-written python GUI (300 epochs, batch size = 4) on 1481 manually annotated images (labelImg, Github) across 14 datasets, ensuring an even spread across the timeframe of each experiment. Object categories labeled with boxes included egg, larvae, mother’s head and mother’s pectoral fins. The trained model (Ext. data Fig. 4) was used separately on every dataset to predict the x and y coordinates of each identified object’s center. Since the position of the shell was fixed for the duration of each experiment, an image of the shell was then broken into three regions of interest (ROI): entrance area, laying chamber and deep chamber, for each experiment using the software labelme (Github). The ROIs were drawn manually (see Fig 1g). Using custom-written python scripts we processed the data, extracting time information and image sequence and then further processed data for the larvae separately to the mothers.

For the larvae, we filtered the preprocessed positional data for only eggs and larvae information and set the threshold of the predicted detection confidence level 0.5 and above. The age of the larvae (dpf) was extracted using the time information and each coordinate was assigned to one of the three ROIs. We used a pivot table to assess the number of larvae in each ROI for each time frame.

From the pivot table we determined the maximum number of larvae detected in each ROI as well as in the whole shell at one time over the experiment. We generated figure 2c and 5a-e, a tile plot, by plotting a vertical line for each frame, separately for each ROI. The color of the line represented the ROI while the alpha component (or transparency) of the color corresponded to the number of larvae in that ROI for the given frame divided by the maximum larvae found in the ROI across the whole experiment, with a high fraction having a higher alpha component. In addition key time points, inputted manually, were plotted onto the figure including the time of parent removal, fresh water provision, food provision and clutch swapping. The emergence time, also plotted on the figure, was automatically detected as the first frame in which two larvae were found in the entrance chamber in the same frame after 5 dpf.

The emergence times were also used to produce Fig. 5f where the experimental runs were grouped according to experimental manipulation and the average emergence time was plotted as a bar graph. Additionally the time larvae move from the laying chamber to the deep chamber was annotated onto the figure and calculated as the last frame in which an egg is detected in the laying chamber.

For the mother, the preprocessed positional data was filtered for prediction confidence above 0.59 for mother’s head and mother’s pectoral fins. To assign the mother’s position to one of the three ROIs, we took the position of the mother’s head and if that was not detected (due to obstruction of the shell whorl), then we took the position of the mother’s pectoral fin (with the highest confidence level) and then assigned her position to one chamber deeper than the location of the pectoral fin, unless it was in the deep chamber already. Similarly to the larvae, we calculated the dpf and used a pivot table to look for the presence of the mother in each ROI for each time frame.

From the pivot table we created Fig. 4a. a tile plot, as described for the larvae, showing the presence of the mother across the different ROIs for each frame, without the alpha component as the mother’s presence was binary. No additional key time points were added to the plots. For each experiment we also plotted the percentage of frames in which the mother visited the shell in the day and the night for each dpf. We collated the visitation data across all experiments, except the cross-fostering experiments, to create a bar graph showing the mean percentage of frames detecting the mothers in the day (Fig. 4b) and the night (Fig. 4c) for each dpf. Data for each dpf was only included when the filming occurred and mother was present in the set up for the full 24 h.

### Mother removals

For the control conditions the fish were left undisturbed for the duration of filming. As the controls, but at 5 or 7 dpf both the mother and father were removed with nets from the tank and the time point noted. Shells with larvae were left untouched.

### Supplying fresh water to the shell

The shell was adapted by drilling a small hole from the outside of the shell into the deep chamber and inserting a tube connector (Harvard, No.72-1475, USA). The tube (0.5 mm inner diameter) attached was then connected to a syringe pump (Aladdin A-1000, Germany) outside the tank with a 50 ml syringe (BD Plastipak, 300865, Germany). Once water flow into the shell was started at 4 dpf (flow rate: 4 ml/hour) it was continuous until the end of the experiment, with the syringe refilled with facility water every 15 hours. Parents were removed at 7 dpf as described above.

### Feeding inside the shell

As the controls, but the same adapted shell for the water flow was used, now connected multiple times on day 8 and 9 post-fertilization to a 10 ml syringe (Norm-ject, Germany) containing 20-30 live artemia, in 5 ml facility water which was perfused into the shell (5 ml/ min).

### Cross fostering experiments

Two females were identified with clutches of eggs two days apart in the breeding facility. When Supplarvae were at 4 and 6 dpf, respectively, they were removed from the mother’s shell into separate 100 mm petri dishes and counted. The experimental tank was divided in two by an opaque barrier, with one filmed open-back shell in each. Larvae clutches were then put into one of the shells with a pipette, ensuring the number of larvae was equal to the size of the smaller of the two clutches. The mothers were added to experimental tanks, receiving the shell containing the clutch from the other mother. Both shells were filmed from the switch point as described above.

### Phototaxis measurement

To test light preference in the larvae, 6 dpf larvae were removed from their mother’s shell in the breeding facility and 4-5 larvae were placed in a custom-made box (30 × 50 mm, 10 mm deep; see Fig. 3a) sitting inside the experimental tank described above, thus exposed to the same water and light conditions described in all previous experiments (see methods: Brood care observations inside the shell). The box was made of clear acrylic with the side walls and half of the lid covered in NIR-penetrable plastic (LUXACRYL-IR,1698, 0.8 mm thick, Germany). This created a box where one half is exposed to, and one half is blocking out, the visible lights in the room, when they are on. After placement, the larvae were allowed to acclimatize before a timelapse (5 images per minute) was started using the same software described earlier (see methods: Brood care observations inside the shell). The time lapse ran from midnight at 7 dpf until 11:59 pm at 11 dpf with a NIR camera (Ximea, MX042RG-CM-X2G2-FF, Germany) with a 50 mm lens (Edmund Optics, Germany), filming from above. The box was not water tight, but larvae could not escape, and the lid, held down by magnets, was flipped once every day by the experimenter between 12-2 pm to change the dark side of the box, during which the filming was temporarily stopped.

To determine the light/dark preferences of the larvae between 7-10 dpf, we used a custom-written python GUI, utilizing labelImg, to manually annotate larvae in 506 images across 6 datasets, ensuring an even spread across the dpf of each experiment. We then used our GUI to train a YOLO model (300 epochs, batch size=4; see Ext. data Fig. 5) which was used separately on the 6 complete datasets to predict the x and y coordinates of each larva’s center point. Using lableme, for each dpf the experiment box was divided into two ROIs, light where the lid was clear and dark where the NIR-penetrable plastic coated the lid. The larvae could be detected equally well on both sides as the NIR camera filmed through the NIR-penetrable plastic.

The position data of all the larvae was processed using a custom made python script for each dpf and all experimental runs. We filtered out predicted detections with a confidence less than 0.25, extracted time information from images, and assigned the dpf and hour of the day for each larva detected. We then assigned the location of each larva to one of the two ROIs and concatenated all dpf files from one experimental run. With a pivot table, we could total the number of larvae in the dark versus the light for each hour of each dpf and find the percentage of larvae in the dark for those time points.

All the experimental runs were collated at this point and we calculated and for figure 3b we plotted the average ± standard error percentage of larvae in the dark for each hour of each dpf. In figure 3c, we compared the average percentage of larvae in the dark during the day pre-emergence (7 and 8 dpf) to that post-emergence (9 and 10 dpf) for each dataset and performed a two-sided Wilcoxon signed-rank test.

Chat GPT v3.5 by OpenAI was used to assist with code writing for some of the analysis.

