## Extended data for "Brood care in shell-dwelling cichlids is timed by independent maternal and larval clocks"

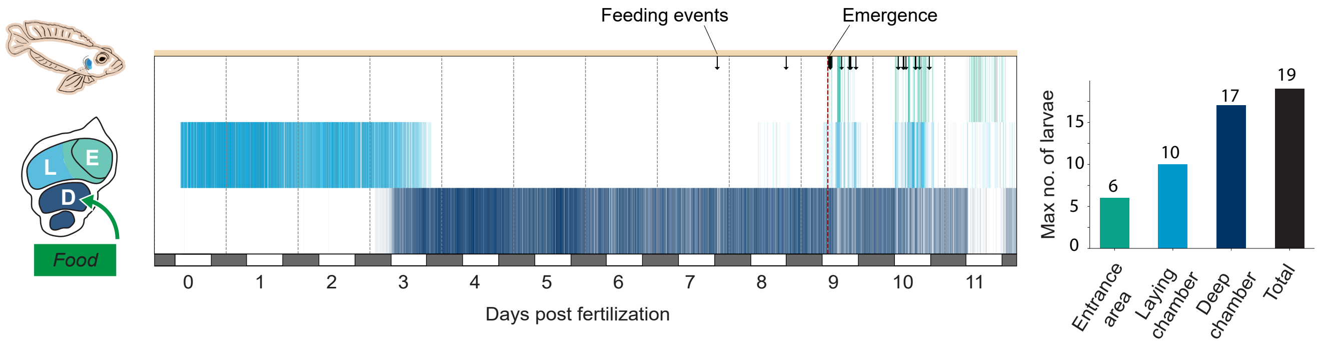


**Extended data figure 1: Larval distribution and emergence during brood care when food was provided into the shell. a)** Larvae distribution across three chambers (E,teal, entrance;L, blue, laying;D, navy, deep) over 11 days post fertilization (dpf) with presence of the mother (orange bar) and emergence time indicated (red line). Black arrows indicate when the experimenter introduced 20-30 artemia in 5 mls tank water into the deep chamber of the shell. White and gray alternating bars under the plots indicate day and night, while vertical dotted gray lines separate each day. Expected emergence on day 9 is indicated by a transparent red bar behind the plots. f) An average of the emergence time of larvae across different manipulations, individual points correspond to each experiment, red dotted line represents the average emergence time in control conditions. **b)** The total number of larvae counted in each chamber and the overall total in the shell. These values were used to normalize the the alpha values in a.

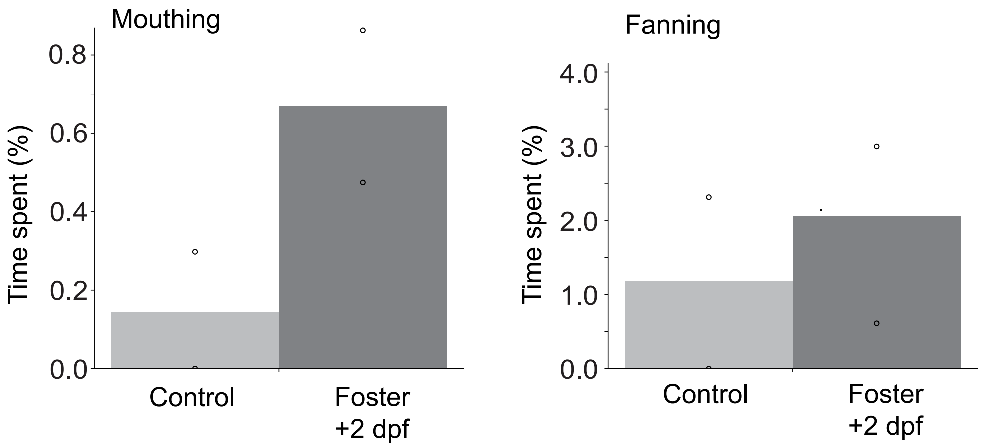


**Extended data figure 2: Control vs foster mother behavior allocations towards 9 dpf lavae.** The percentage time control mothers (with biological offspring) compared to mothers fostering larvae 2 days older than her original clutch, spend mouthing (n=2) **(a)** or fanning (n=2) **(b)** the 9 dpf larvae during 20 min video recordings at noon. Bars graph represents the mean percentage across mothers recorded and circles are the values for the individual mothers.

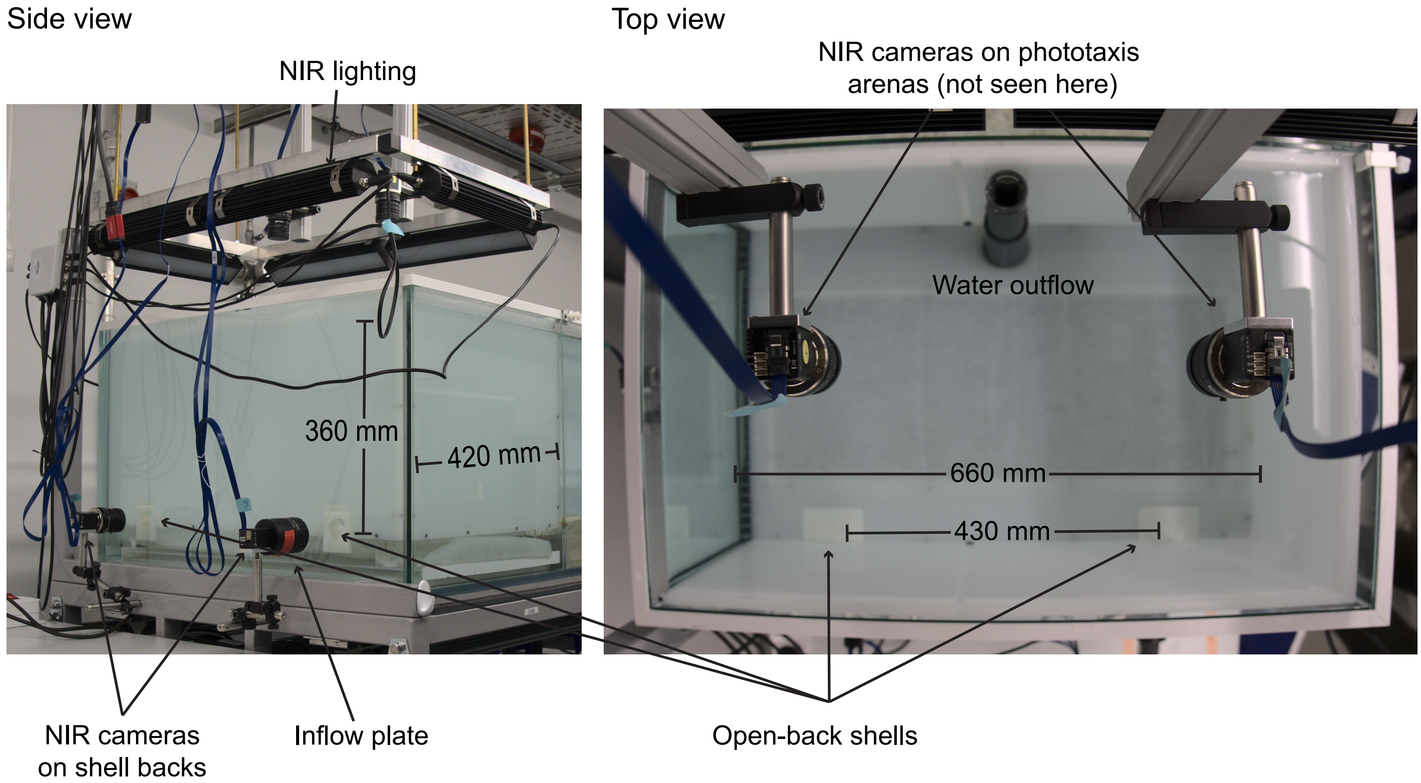


**Extended data figure 3: Self-constructed 100 l tank used for all behaviour experiments.** The design allowed for the insertion of two open-back shells, for the male and female *Lamprologus ocellatus,* into the light-absorbing acrylic tank insert. Under the tank insert was an inflow plate with an even distribution and an outflow tube from the surface on the opposite side to the shells. The shell backs were imaged by near-infrared (NIR) cameras for observations of the larvae and mother inside the shell. The top view NIR cameras were used in the phototaxis experiments, where the small box was placed under the camera on the floor of the experimental tank. The tank was lit by 6 NIR lights from above and two NIR lights (not imaged) stood next to the camera filming the shell backs to flood NIR light into the shell. Ambient room light on a 13:11 light/dark cycle was blocked from entering the shell from the back with a strip of NIR-penetrable black plastic (not imaged).

**Extended data table 1: Larvae behavioral repitoire scored in the BORIS software.** The behavior code, type of event and the description of the behavior used when scoring videos in BORIS

| Behavior code | Type | Description |
|---|---|---|
| Head attached | State event | Larva head attached to shell wall |
| Resting on floor | State event | Head not attached but larva is weighed down by yolk, lying on shell floor |
| Wriggling | State event | Larva performs high frequency tail beating |
| Swimming | State event | When larva are actively or passively displaced within the shell |

**Extended data table 2: Mother behavioral repitoire scored in the BORIS software.** The behavior code, type of event and the description of the behavior used when scoring videos in BORIS

| Behavior code | Type | Description |
|---|---|---|
| Mouthing | State event | Mother actively engages with eggs or larvae using her lips or picking up individuals and circulating them in her buccal cavity |
| Fanning | State event | Mother rapidly moves her anal and caudal fin to facilitate water exchange inside the shell |

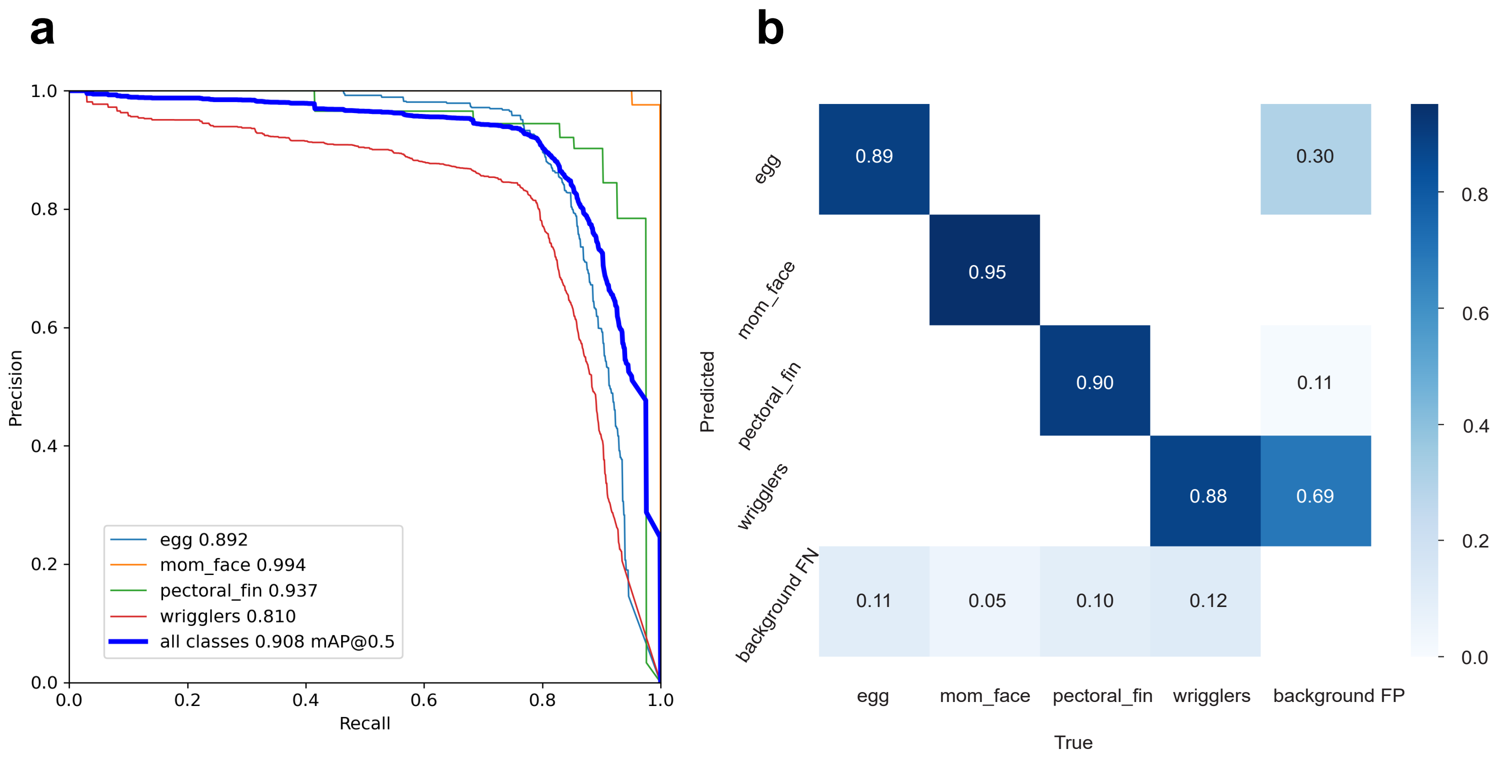


**Extended data figure 3: YOLO neural network evalutation metrics for object detection inside the shell.** a) The precision recall curves and average precision values for each object class (blue, mom; orange, mom_face; green, pectoral_fin and red, wrigglers) and the combination of all classes (navy), with an mean avergae precision (mAP) at 0.5. b) The confusion matrix for the four above mentioned classes and the background.

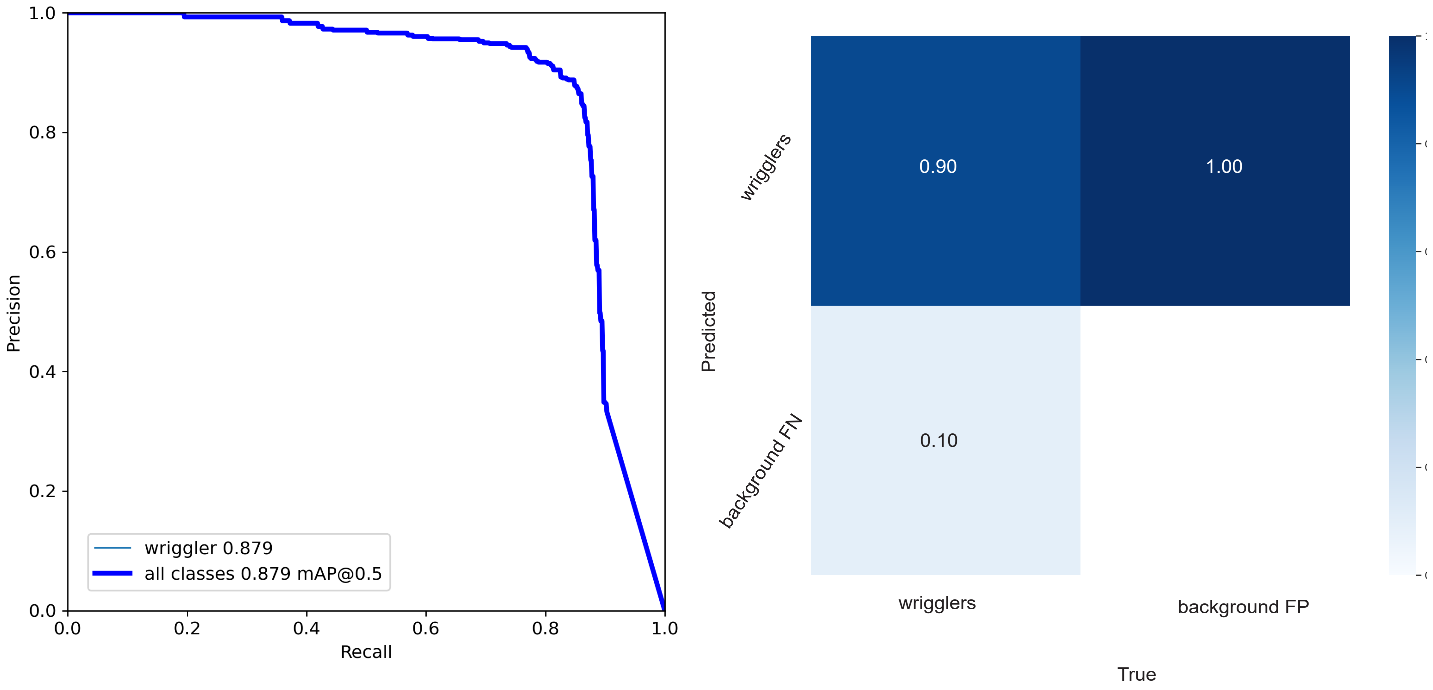


**Extended data figure 4: YOLO neural network evalutation metrics for object detection in phototaxis assay.** a) The precision recall curves and average precision values for wrigglers, with an mean avergae precision (mAP) at 0.5. b) The confusion matrix for the wriggler dectections and the background.
